# Local Homogeneity of Tonotopic Organization in the Primary Auditory Cortex of Marmosets

**DOI:** 10.1101/398677

**Authors:** Huan-huan Zeng, Jun-feng Huang, Ming Chen, Yun-qing Wen, Zhi-ming Shen, Mu-ming Poo

## Abstract

Marmoset has emerged as a useful non-human primate species for studying the brain structure and function. Previous studies on the mouse primary auditory cortex (A1) showed that neurons with preferential frequency tuning responses are mixed within local cortical regions, despite a large-scale tonotopic organization. Here we found that frequency tuning properties of marmoset A1 neurons are highly uniform within local cortical regions. We first defined tonotopic map of A1 using intrinsic optical imaging, and then used *in vivo* two-photon calcium imaging of large neuronal populations to examine the tonotopic preference at the single-cell level. We found that tuning preferences of layer 2/3 neurons were highly homogeneous over hundreds of micrometers in both horizontal and vertical directions. Thus, marmoset A1 neurons are distributed in a tonotopic manner at both macro- and microscopic levels. Such organization is likely to be important for the organization of auditory circuits in the primate brain.

## Introduction

Topographic cortical maps are essential for sensory perception and animal behaviors. In the auditory system, the most prominent topographic feature is tonotopic organization, in which adjacent cortical regions showed preferential responses to pure tones of nearby frequencies. In many species, *in vivo* electrophysiological recordings and imaging techniques have characterized the global tonotopic organization of the auditory cortex, revealing its division into separate fields and distinct tonotopic maps within each field^1–8^. Although large-scale imaging and recording methods often yield global tonotopic maps, recent studies using two-photon calcium imaging to monitor the activity of individual neurons in mice showed that local populations of A1 neurons were highly heterogeneous in their frequency-tuning properties, although macroscopic imaging using intrinsic optical imaging over the entire auditory cortex showed an overall tonotopic organization^9–11^. Thus, macroscopic tonotopic map may reflect an averaged frequency preference for large populations of neurons with heterogeneous tuning properties^12–14^.

Marmoset is a species of new world monkeys known to be highly vocal and social^14^. It has a neocortex much closer to humans than the commonly used rodent models. The spatial distribution of cortical neurons with different frequency preferences is important for the organization of neural circuits that processes auditory signals, such as natural sounds comprising complex mixture of frequencies. Thus, it is important to determine whether the local heterogeneity in neuronal frequency tuning found for mouse A1 neurons is a general property of mammalian auditory cortices, or alternatively, a property more unique to rodent brains. To address this issue, we used *in vivo* two-photon imaging to monitor pure tone-evoked responses of a few hundred A1 neurons simultaneously in anesthetized common marmoset (*Callithrix jacchus*). We found that A1 neurons within distances of a few hundred micrometers in both horizontal and vertical directions were highly homogeneous in their frequency tuning properties. We also showed that this tonotopic organization in marmoset A1 is distinctly different from that found in rat A1 by the same imaging method. Such micro-architecture of the auditory cortex may be important for efficient coding of natural sounds in highly vocal animals such as marmosets.

## Results

### *In vivo* two-photon calcium imaging in marmoset A1

To facilitate identification of various areas of marmoset auditory cortices, we performed intrinsic optical imaging^15^ in anesthetized marmosets (see Methods). We observed that pure tone stimuli could evoke intrinsic optical signals in three primary auditory regions, previously defined by electrophysiological recording and anatomical studies^16–19^ as the primary (“A1”), rostral field (“R”) and rostro-temporal field (”RT”) of the auditory cortex (**Fig. 1a-d** and **Supplementary Fig. 1**). For A1 lying on the ventral bank of lateral sulcus (LS), we found that gradual increments of sound stimuli from low- to high-frequencies evoked responses in adjacent areas along LS from the antero-ventral region to postero-dorsal region of A1 (**Fig. 1a-c).** The overall tonotopic frequency-angle map is shown for one example imaging plane in **Fig. 1d**, with different colors indicating frequency preferences. Similar map was observed in two other marmosets (**Supplementary Fig. 1**). This finding is consistent with previous results obtained by electrophysiological and optical imaging approaches^7,8^.

**Figure 1.**
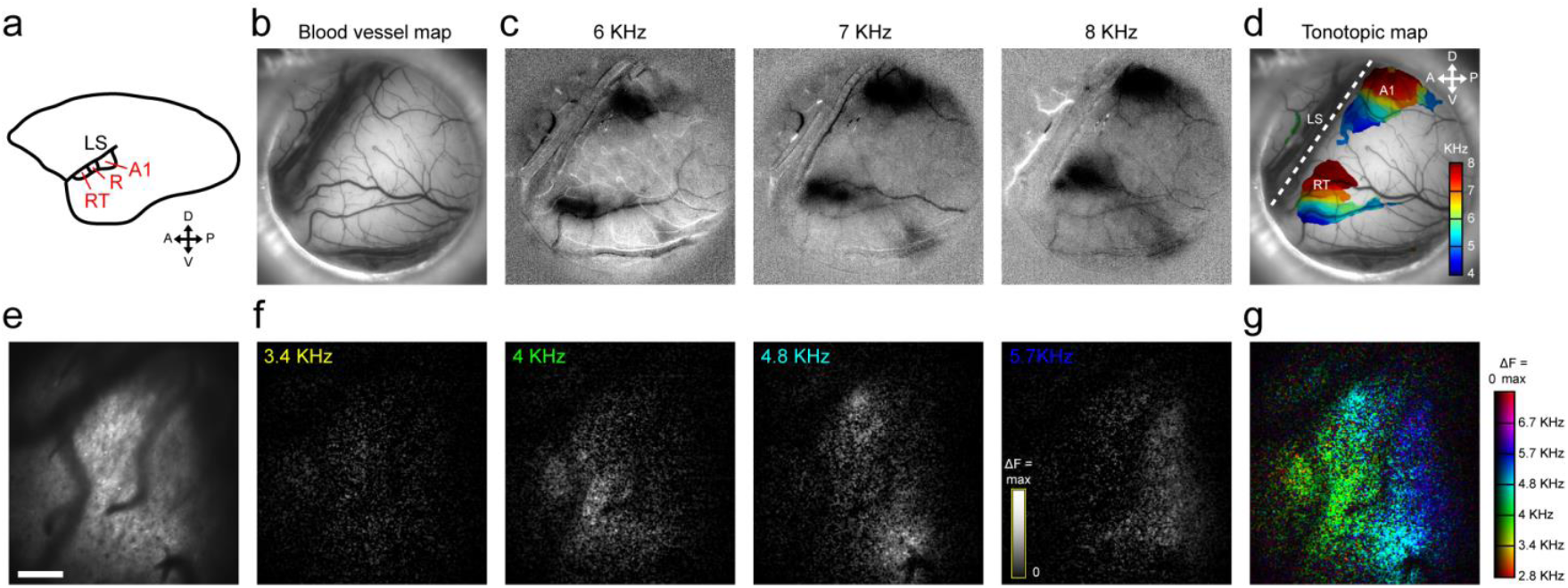
Large-scale imaging of neuronal responses in the marmoset auditory cortex. (**a**) Schematic diagram showing various regions of the marmoset auditory cortex. LS, lateral sulcus; A1: primary auditory cortex; RT, rostro-temporal field; R, rostral field. (**b**) Surface blood vessel map within the imaging window. (**c**) Intrinsic optical signals evoked by pure tone stimuli at 6, 7, 8 KHz, respectively. Dark areas, activated regions. (**d**) Summary of tonotopic maps (in the left hemisphere) revealed by intrinsic optical imaging using pure tone stimuli, color-coded for five discrete frequencies (4 to 8 kHz). Note the gradual shift of the responses along LS. (**e**) Imaging plane obtained using a 16X objective (with 2-μm pixel-1). (**f**) Pure-tone respobnses (ΔF) for 4 different frequencies. Each image represents the average of five repeats. (**g**) Composite frequency-preference map for the imaging plane as in **e**. Colors indicate the preferred frequencies; with the brightness indicating the magnitude of the frequency selectivity. Scale bars:1 mm in **b-d**; 200 μm in **e-g**.

Following obtaining the overall frequency preference map of A1 with imaging of intrinsic optical signals, we further performed *in vivo* two-photon calcium imaging at selected local A1 regions by bulk-loading with the fluorescent calcium indicator Cal-520 AM^20^. Loading of the indicator reached an apparent plateau at ∼60 min after local perfusion, when hundreds of fluorescent cells could be detected at imaging depths that cover the major part of the cortical layers 2/3. Large-scale imaging over area of about 1 mm × 1 mm with a 16X objective lens allowed us to measure global frequency tuning of A1 sub-regions, using a method similar to those reported previously for studying orientation and spatial frequency tuning in the primary visual cortex^21,22^. By measuring the fluorescence changes (ΔF) of all pixels within the imaging plane in response to pure tones at six discrete frequencies from 2.8 to 6.7 KHz, we found that activated regions were organized tonotopically (**Fig. 1e-g**). Thus, large-scale imaging using both intrinsic and Ca^2+^ signals revealed similar tonotopic organization of marmoset A1, confirming previous electrophysiological findings.

### Clustered distribution of A1 neurons with similar frequency tuning

Two-photon imaging of Ca^2+^ fluorescence signals at a higher resolution (with 40X objective, ∼0.6μm/pixel) allowed us to monitor pure tone-evoked Cal-520 AM fluorescence changes in individual A1 cells within a given focal plane (see Methods). We found that most cells exhibited maximal responses at a specific frequency (defined as the best frequency, “BF”). The percentage of responsive cells among all fluorescently loaded cells was 48 ± 6 % (SE, n=5 fields), and cells with similar BFs were found to localize together within the imaged field. In the example imaging field shown in **Fig. 2a**, 10 cells sampled within a distance of about 180 μm all showed the BF at 8 kHz. When fluorescence signals from all 68 responsive cells were measured over the entire imaged field, for pure-tone stimuli over discrete incremental frequencies from 0.5 to 32 kHz, we found most cells had BFs centered around 8 kHz (**Fig. 2b**). This frequency preference was also shown by the average ΔF/F tuning profile for all 68 cells (**Fig. 2b**, **top inset**). This homogeneity of BFs could also be visualized by the tonotopic map with BFs coded by discrete colors (**Fig. 2c**). As summarized by the histograms for the percentages of cells exhibiting different BFs, we observed such local uniformity of BFs for 5 different imaging planes (ranging from 180 to 520 μm in width) recorded from 3 marmosets over the frequency range from 1 to 9.5 kHz (**Fig. 2d**). Such homogeneity of tonotopic properties within each imaged area is in sharp contrast to that found in mice using Ca^2+^ imaging methods ^9,10^, where cells within 50-100 μm distances showed much larger variability in BFs (up to four octaves).

**Figure 2.**
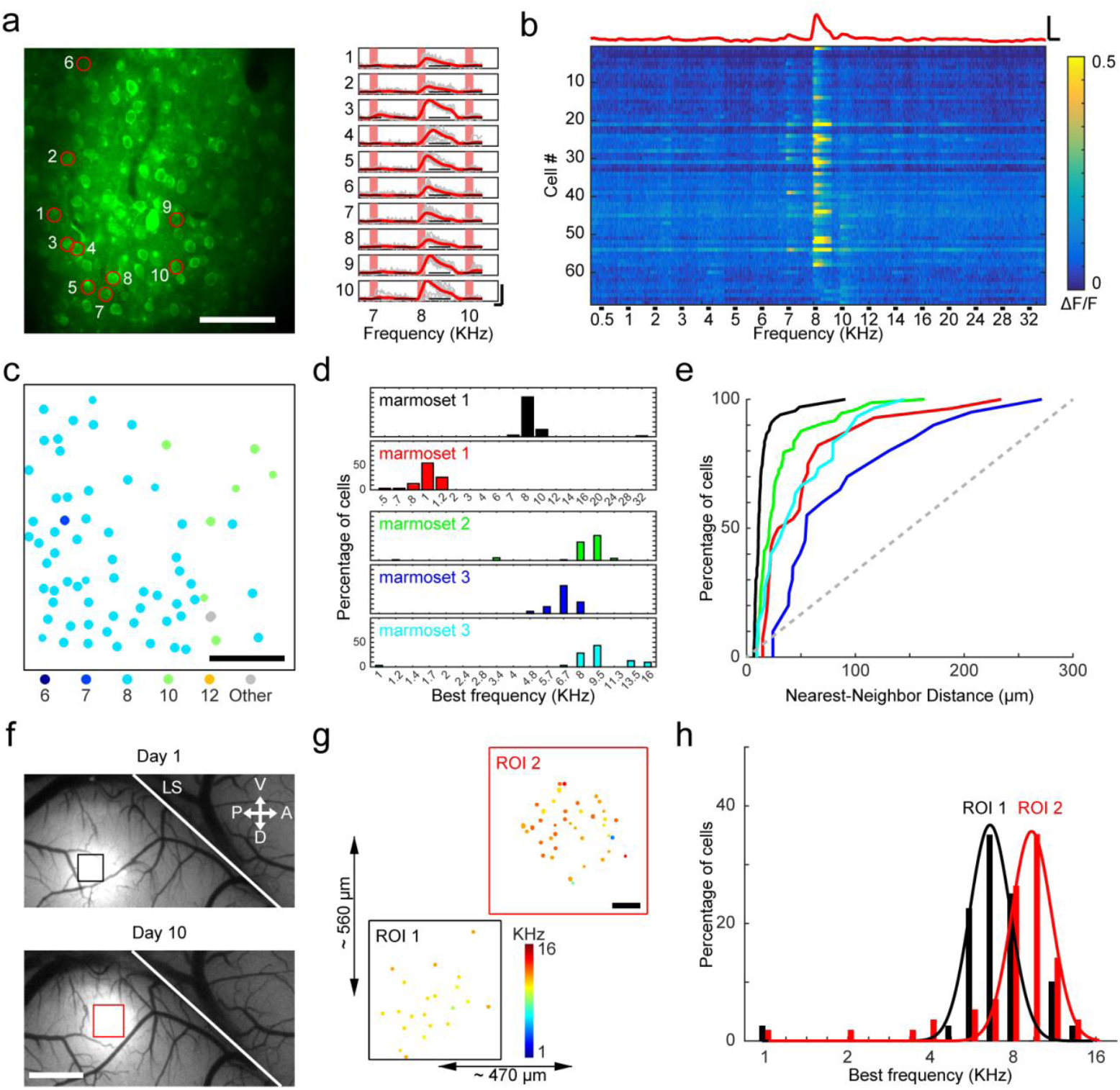
Marmoset A1 cells with similar frequency preference were highly clustered within local cortical regions. (**a**) **Left**: *In vivo* two-photon image of Ca^2+^ fluorescence at a focal plane, at ∼150 μm below the pia surface. Bar, 50 μm. **Right**: Single trials (gray lines, n = 8) and mean (red line) fluorescence changes (ΔF/F) evoked by pure tone stimuli (7-10 KHz) at ten sampled cells (marked by red circles on the **Left**). Scale bar, 0.2s and 20% ΔF/F. (**b**) ΔF/F with time for 68 pure tone-selective cells during sequential step-wise application of discrete pure tones from 0.5 to 32 kHz (black bar, tone duration of 0.2 sec), with the magnitude of ΔF/F color-coded by the continuous scale shown on the right. Red trace on top shows the averaged ΔF/F for all 68 cells. Note that the best frequency (BF) for the vast majority of cells was located at 8 KHz. Scale bar, 0.5s and 20% ΔF/F. (**c**) Spatial distribution of all 68 cells shown in **a**, with their BFs (KHz) color-coded by the scheme shown on the bottom. Bar, 50 μm. (**d**) The percentage of cells with different BFs observed in five imaged planes from 3 marmosets. Top panel corresponds to the example imaging plane shown in **a-c**. (**e**) Cumulative percentage plot for the distribution of nearest-neighbor distances for all cells in each of five different imaging planes as that in **d** (same color coding. **Black**: data for 68 cells shown in **a-c**. **Colored**: data collected from 4 other imaging planes observed in 3 different marmosets. Gray diagonal line depicts uniform distribution). All curves were significantly different from the diagonal line (P < 0.001 for all curves, Kolmogorov-Smirnov test). **(f)** Fluorescence images of two adjacent regions in marmoset A1 after Cal-520 loading of the same marmoset in two experiments performed 9 days apart. Bar, 1 mm. (**g**) Distribution of cells with different BFs, in the two regions of interest (ROIs) boxed in **f**. Bar, 100 μm. (**h**) Histograms of BF distributions for all tone-selective cells recorded in the two ROIs. (n = 39 in ROI 1, n = 57 in ROI 2).

We also quantified this clustering of cells with the same BFs by calculating the nearest-neighbor distance, defined as the distance of the nearest cell showing the same BF, for all responsive cells in the imaging plane (**Fig. 2e)**. The cumulative percentage plot of the distribution of nearest-neighbor distances, of all five imaged planes showed steep slopes at small distances, with median distance (50%) of 30 ± 7 μm (SE, n = 5). These distributions were significantly different from the random uniform distribution of near-neighbor distances (P<0.001, Kolmogorov-Smirnov test), consistent with the homogeneity of BFs within the imaged plane.

In the experiments above, we have shown a tonotopic organization of A1 at the millimeter scale using imaging of intrinsic optical signals and local homogeneous distribution of cells with the same BFs. To further confirm that the local homogeneous tonotopy occurred at different A1 regions in the same marmoset, we performed measurements on two adjacent A1 areas which were separated by about 500 μm (**Fig. 2f**), in two separate experiments 9 days apart. We observed a clear difference in the dominant BFs of 6.7 and 9.5 kHz in the two regions respectively, as shown by the BF distribution histograms (**Fig. 2h**). These experiments also showed the feasibility of repeated long-term two-photon imaging of cortical neuronal populations over many days in marmosets.

### Tonotopic homogeneity in A1 is sound intensity-invariant

We next examined whether the BF of the same A1 cell depends on the intensity of the sound stimuli. Using pure tones of three sound intensities (60, 70 and 80 dB), we found that many A1 cells exhibited higher responses as the sound intensity was increased, while others showed similar responses at all three intensities, and a few showed reduced responses at higher intensities (**Fig. 3e and f**). Despite the variation in the dependence on sound intensity, the vast majority of A1 cells within the same imaging plane were narrowly tuned to the same frequency at all three sound levels, as shown by the cell-based tonotopic maps (**Fig. 3a**, BFs around 8 KHz). As shown for the example cell in **Fig. 3b**, similar averaged response profiles and BF distributions were observed for the responses at three different sound intensities (all peaked at 8 KHz, **Fig. 3c**). This is largely consistent with the previous finding using electrophysiological recording that sound-evoked responses of awake marmoset A1 neurons is sound level-invariant^23^. Further analysis indicates that the distributions of nearest-neighbor distances of the same BFs were all significantly different from the distribution expected for uniform distribution (**Fig. 3d**, *P*<0.001, Kolmogorov-Smirnov test).

**Figure 3.**
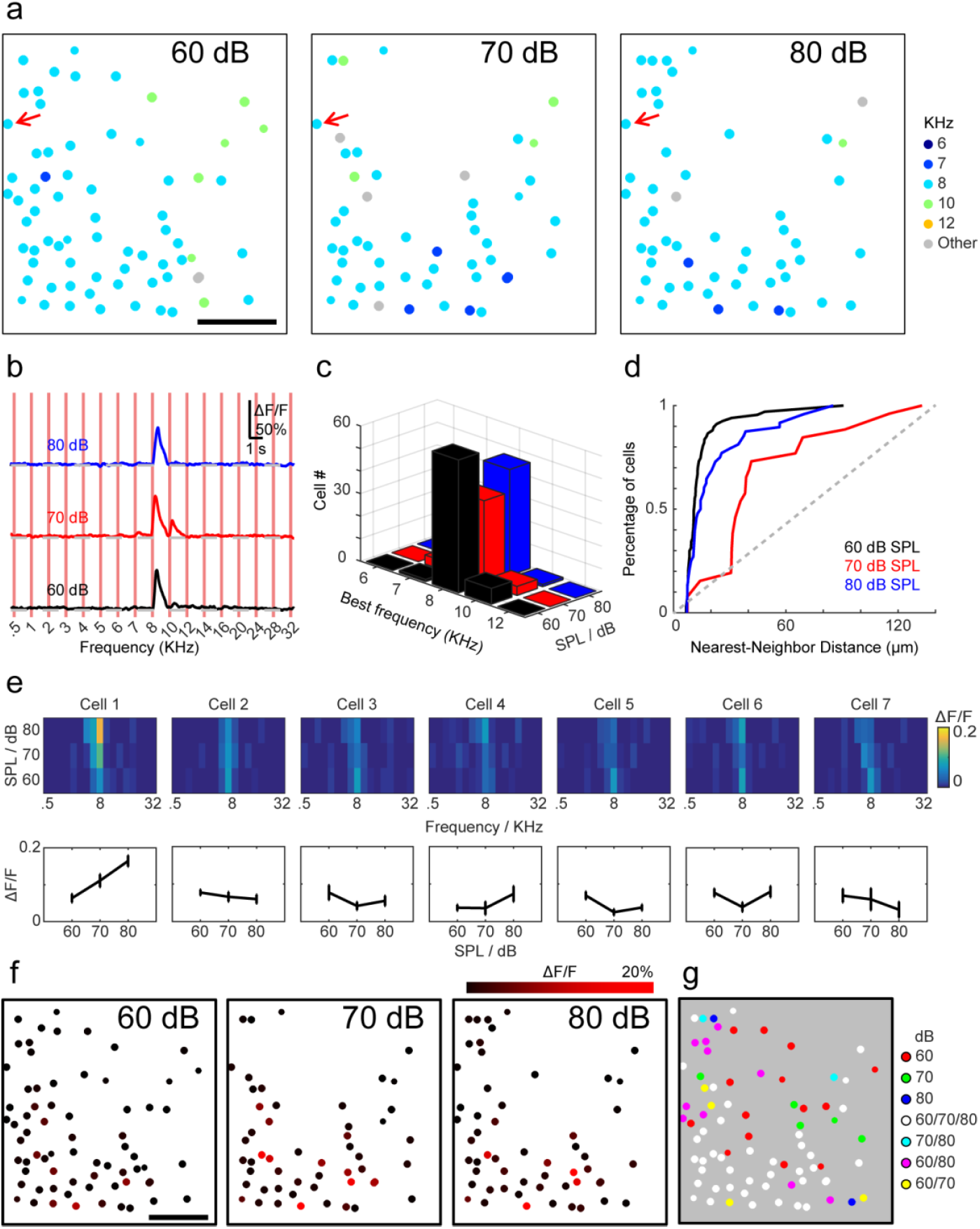
Local homogeneity in frequency tuning did not depend on the sound intensity. (**a**) Sound-evoked responses of the same population of A1 cells at three different sound intensities (60, 70, 80 dB), with BFs of the cells color-coded. Data were collected for the same imaging plane as that shown in Fig. 2. Bar, 50 μm. (**b**) The averaged frequency tuning profiles derived from an example cell (indicated by red arrows in **a**) at three different sound intensities (60, 70, 80 dB). (**c**) Total number of cells showing different BFs for three different sound intensities. (**d**) Cumulative percentage plot for the distribution of the nearest-neighbor distance for all cells at three different sound intensities. All curves were significantly different from the diagonal line (P < 0.001 for all curves, Kolmogorov-Smirnov test). (**e**) Tonal receptive fields (TRFs) of 7 example cells from the same imaging plane in **a** are shown. Plots below depict maximum mean fluorescence changes to pure tone stimuli with increasing intensities, showing diverse changes in cellular responses at three different sound intensities. (**f**) Cells in one imaging plane (as in **a**) that responded to the pure tone at a particular sound intensity. The brightness of the dots indicates the response magnitude (maximum mean ΔF/F, color-coded by the scale bar). Bar, 50 μm. (**g**) Color-coded intensity map derived by combining the three distribution plots shown in **f**. Colors of cells indicate whether the cells showed responses to one, two, or three intensities. Note that most cells were activated at all three intensities (white dots).

Previous single-unit recordings from A1 of several species have suggested a patchy organization for intensity tuning along the iso-frequency axis^24–26^. Thus we have examined the existence of clustering of cells with the same BFs based on the preferred intensity, by the distribution of cells with different BFs in the imaging plane (**Fig. 3f**). Although some cells were best activated at different intensities, the number of cells activated were similar, with 68, 51, and 54 (out of 109) activated at three intensities tested, respectively. We noted that most cells could be activated by all three intensities (**Fig. 3g**; 40 white dots) and a small fraction of cells showed intensity selectivity (**Fig. 3g**; 14 red dots, 5 green dots, 2 blue dots). However, there was neither apparent clustering of cells that respond selectively to one particular intensity nor an apparent gradient of best intensity across the iso-frequency axis.

### Vertical organization of A1 tonotopic maps at superficial cortical layers

A general organization principle in many sensory cortices is that neurons for processing similar functional features are organized into vertical columns^27,28^. We thus further examine whether tonotopic maps are also homogeneous along the vertical axis of A1. Due to the technical limitation of our two-photon Ca^2+^ imaging method for the marmoset, we were only able to address this issue by exploring the tonotopic properties of A1 neurons with a depth up to 370 μm from the pial surface, covering the major part of layer 2/3. **Fig. 4a** shows the spatial distribution of A1 cells, color-coded with their BFs in the example experiment shown in **Fig. 2**. Different groups of cells at five different focal depths (140, 170, 200, 230, and 260 μm) were all found to exhibit BFs predominately at 8 or 10 KHz, with clear clustering of cells of similar BFs.

**Figure 4.**
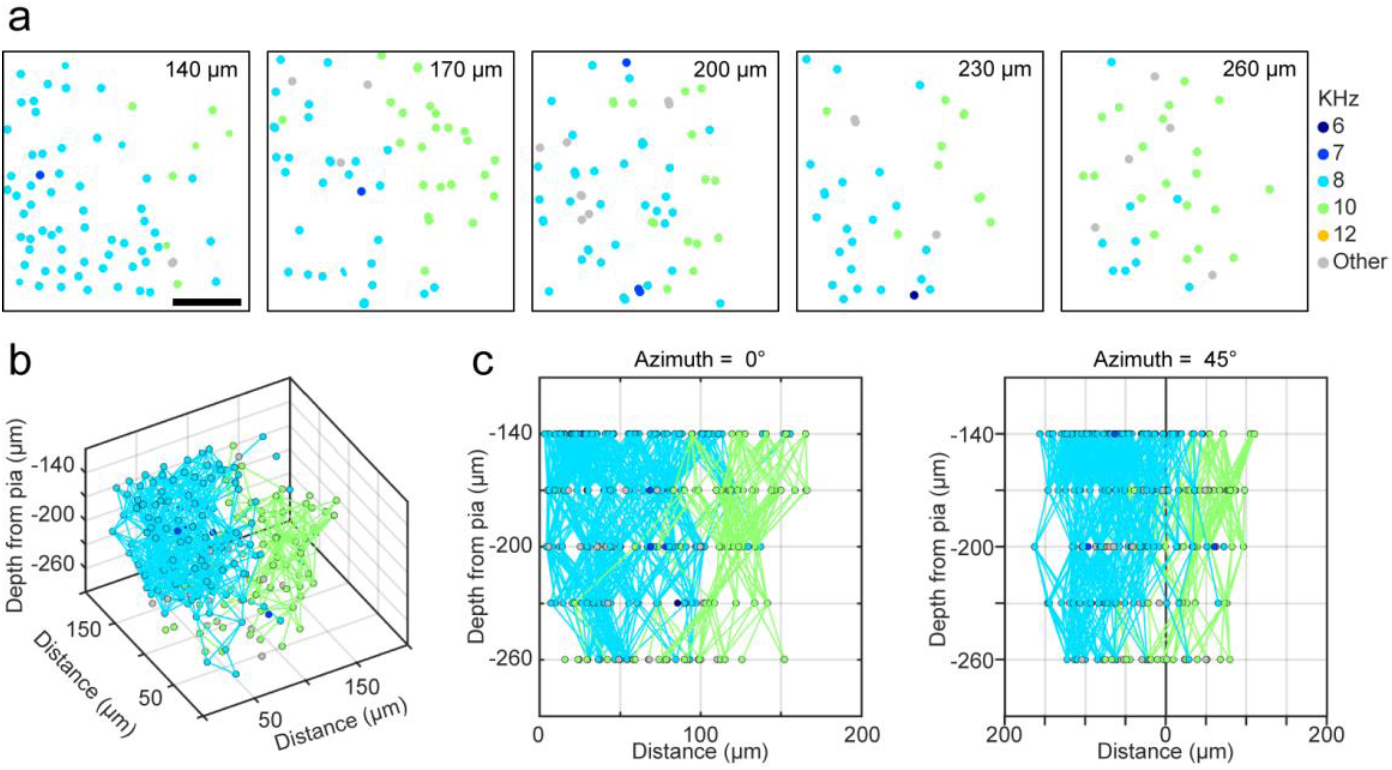
**(a)** Spatial distribution of A1 cells with distinct BFs at five different cortical depths from 140 to 260 μm, with the BF of each cell color-coded by the scheme shown on the right. **(b)** 3D composite plot of the distribution of cells with the same BFs, with cells of the same BFs in different depths within 50 μm linked by lines of the same color as that coding BFs. (**c**) Projected 2D maps of the composite plot onto two planes at azimuth angles of 0 and 45 degree, showing vertical alignment of cells with the same BFs at different depths. Bar 50 μm.

Composite 3-D plot (cell pairs sharing the same BFs from adjacent imaging planes with a distance < 50 μm were connected by lines) of the cell distribution showed that clusters of cells with the same BFs are well aligned vertically among imaged planes, although there appeared to be more cells with BFs deviated from 10 kHz at deeper cortical regions (**Fig. 4b**). In a separate experiment on a different marmoset, we imaged 5 planes over the cortical depths between 250-370 μm, similar clustering and vertical alignment of cells of the same BFs were also observed. These findings support the existence of columnar structure of A1 tonotopic maps at the microscopic level.

### Local tonotopic organization is more heterogeneous in Rat A1

The above studies showed that the tonotopic organization in the marmoset A1 is highly homogeneous. Given the previous findings showing marked local heterogeneity in the frequency tuning of mouse A1 neurons, we further examine the tonotopic organization of rat A1 neurons using the same Cal-520 imaging method as that used in the marmoset studies above. We found that, for anesthetized SD rats (∼300 g), A1 cells within a local area (∼307×324 μm) in general showed BFs from a low (2 KHz) to high frequencies (32 KHz), with a range that covered 4 octaves (**Fig. 5a**). By contrast, over a larger size of imaging area (∼524×554 μm) of marmoset A1, the BFs ranged less than 2 octaves **(Fig. 5b)**. Rough inspection of the local tonotopic maps of rat A1 revealed many regions exhibited “salt and pepper” distributions of BFs (see box 1, **Fig. 5a**), although small regions with clustered distribution of similar BFs could also be found (see box 2, **Fig. 5a**).

**Figure 5.**
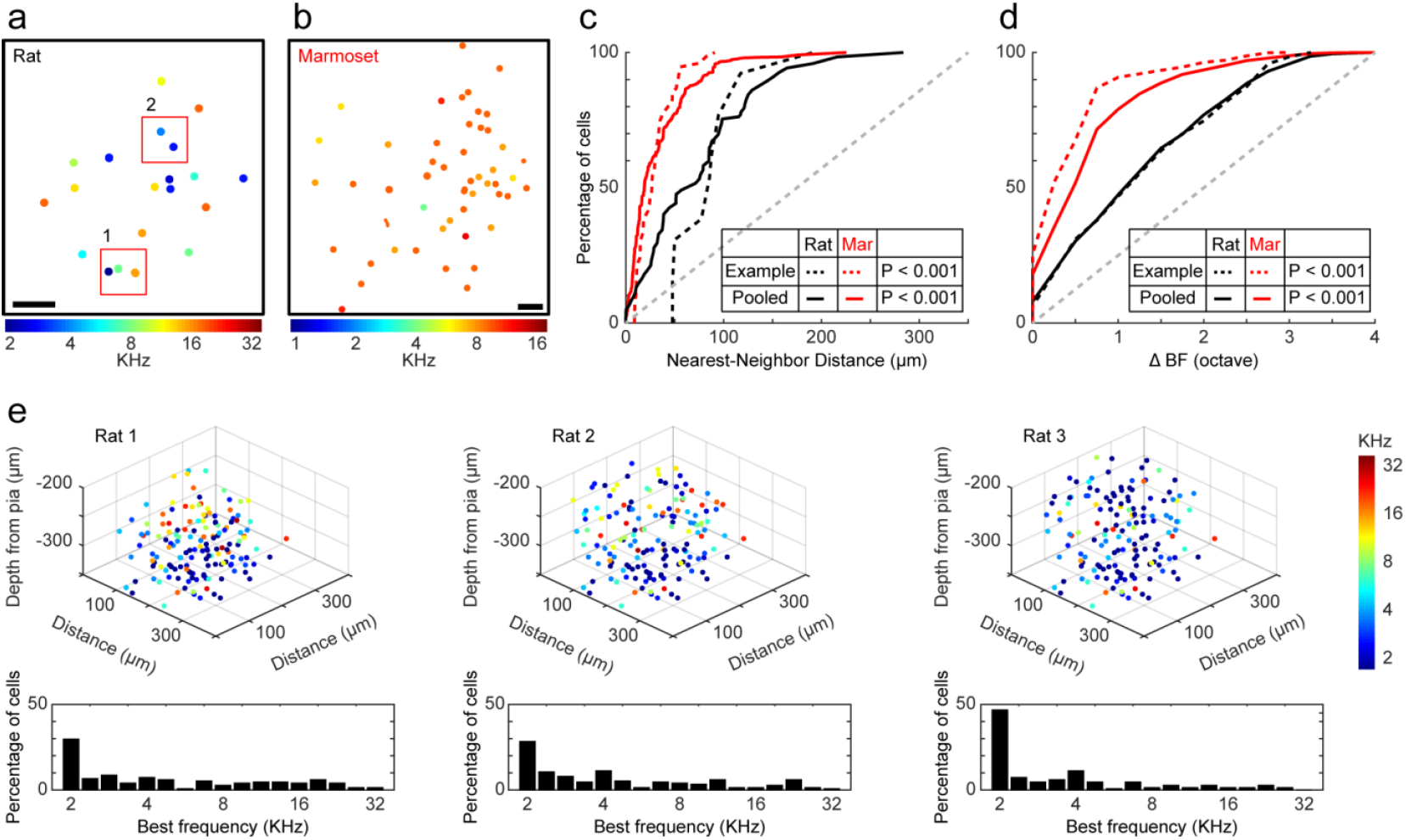
Local tonotopic organization in rat A1 is more heterogeneous than that in marmoset A1. (**a**) Spatial distribution of all tone-selective cells in an example imaging plane in the rat A1, with their BFs (KHz) color-coded by the scheme below. Red boxes indicate 2 sub-regions with “clustered” or “salt and pepper” distribution. Bar, 50 μm. (**b**) Spatial distribution of all tone-selective cells in an example imaging plane from a marmoset. Note the uniform distribution of BFs. **(c**, **d)** Cumulative percentage plots for the distribution of nearest-neighbor distances of the same BF for all cells (in c) and for the distribution of ΔBF for all cell pairs within the same imaging plane (in d), respectively. Red, marmoset; black, rat. **Dashed lines**, data for the example imaging plane as shown in a and b; **solid lines**, pooled data from 9 imaging planes from 2 rats and 11 imaging planes from 2 marmosets. **Diagonal dashed line**, random (uniform) distribution. All curves were significantly different from the diagonal line (P < 0.001 for all curves, Kolmogorov-Smirnov test). Curves for rats and marmosets were significantly different (P < 0.001 for all curves, Kolmogorov-Smirnov test). (**e**) Spatial distribution of A1 cells with various BFs at different cortical depths from 200 to 350 μm. Data from three different rats. The percentages of cells with different BFs in each imaging region are shown in the bottom histograms. Note that in all three rats, similar A1 regions (with dominant BF of 2 KHz) were imaged.

Two types of quantitative analyses were formed to examine the uniformity in the BF distribution within A1. First, we measured the nearest neighbor distance of the cell with the same BFs for all responsive A1 cells monitored in the same image plane (as that done earlier for marmoset, **Fig. 2**). This analysis showed that the distribution of nearest neighbor distances in rat A1 was much closer to the random distribution, as shown by the cumulative percentage curves for the example cases and averaged data from two rats (9 imaging planes) and two marmosets (11 imaging planes) (**Fig. 5c,** 9 imaging planes from 2 rats and 11 imaging planes from 2 marmosets). In the second analysis, we measured the ΔBF for all cell pairs within the imaged plane, and plotted the cumulative percentage curves for both rat and marmoset A1, based on the same data set as above. This analysis showed that the distributions of ΔBF for rat A1 was significantly different from those of marmoset (P<0.001, Kolmogorov-Smirnov test), much closer to the random distribution (diagonal line), although both marmoset and rat distributions were significantly different from random (P<0.001, Kolmogorov-Smirnov test) (**Fig. 5d**). Taken together, our results indicate that, over distances of hundreds of micrometers, local A1 tonotopic map in rats exhibited a much higher heterogeneity than that in marmosets.

To examine whether the non-uniform local tonotopic organization exists along the vertical axis of rat A1, we have also imaged the cellular tuning responses within a local region (∼307×324 μm) at different cortical depths (210-350 μm from pia surface) in three rats examined in this study. We found that for a similar imaging area in A1, cells with diverse BFs were distributed in similar ‘’salt and pepper’’ distribution in all three rats and there was a dominant population of cells with the same BF (2 KHz) in all three rats (**Fig. 5e**). These findings indicates there was reproducible tonotopic map among different rats, and the existence of predominant BF in local A1 region could account for the global tonotopic property in rodents, despite the presence of local heterogeneity in BFs.

### Tonotopic responses of A1 cells using genetically encoded Ca^2+^ indicators

The above results were obtained by using fluorescent Ca^2+^ indicator Cal-520 that were acutely loaded into neurons. It is known that due to differential Ca^2+^ affinities of different indicators, the recorded neuronal activities may be biased by the extent of subthreshold responses included in the fluorescence signal^10,29^. Although Cal-520 has a *K*_*d*_ (320 nM) similar to that of Fluo-4 (350 nM), which is known to respond only to suprathreshold activities^10^, we decided to perform imaging experiments using a genetically encoded ultrasensitive fluorescent protein GCaMP6f (ref. 30), which can reliably detect single action potentials in marmoset neurons^31,32^.

Marmosets were injected with AAV vectors encoding human synapsin promoter-driven GCaMP6f in A1 at 3 weeks prior to the imaging experiment (see methods). In a large focal plane (∼520×550 μm, **Fig. 6a**), A1 cells expressing GCaMP6f showed robust responses to single pure-tone stimuli, as shown by the tone-evoked fluorescence changes in 4 example cells at different sound frequencies (**Fig. 6b**) as well as by the heat map of florescent changes recorded from 20 responsive cells in one imaging plane (**Fig. 6c**). These results indicate that the BFs were homogeneous over large distances in the imaging plane. The spatial distribution of cells with different BFs also showed homogeneous distribution of BFs within local regions of A1, as shown in **Fig. 6d**. In constructing this distribution map, we have combined the data for cells activated by sound stimuli at three different intensities, due to the relatively low number of cells expressing GCaMP6f, as compared to that observed with Cal-520 loading. We also examined the tonotopic responses of A1 cells evoked by sounds at various intensities and found the BFs were largely invariant over three levels of sound intensities (60, 70, and 80 dB). When imaging planes at three different cortical depths (180, 220 and 250 um) were examined for their tonotopic organization, we found again similar clustering of cells with the same BFs at three different depths. These results were summarized by the 3D plots, with cell pairs sharing similar BFs from adjacent imaging planes within a distance < 100 μm were connected by lines (**Fig. 6e**). Taken together, our experiments using GCaMP6f have largely confirmed the findings using Cal-520, showing both horizontal and vertical local homogeneity in sound frequency preference and intensity invariance of tonotopic properties in marmoset A1 neurons.

**Figure 6.**
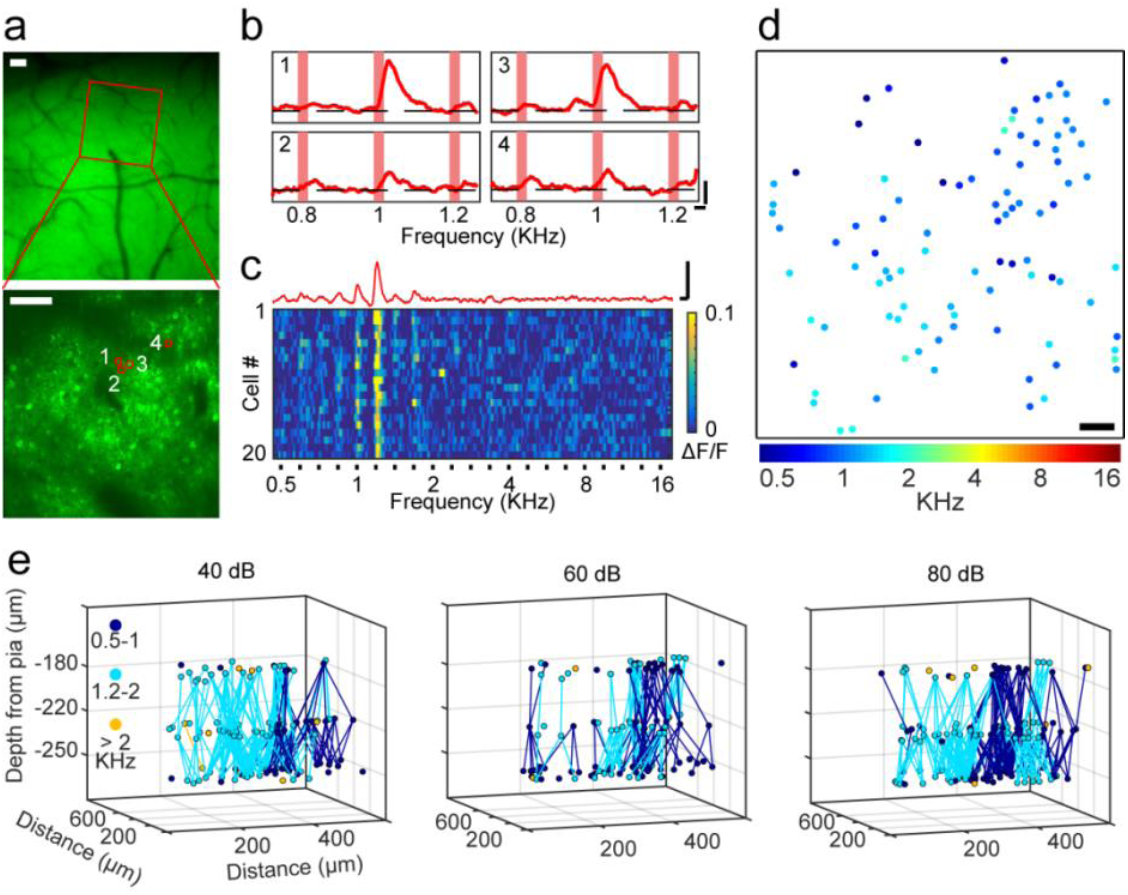
Two-photon imaging of marmoset A1 using genetically encoded Ca indicator GCaMP6f. (**a**) **Top**: Fluorescence image of a region in marmoset A1, 3 weeks after AAV-GCaMP6f injection. **Bottom**: Two-photon fluorescence image of a ∼520×550 μm imaging area in A1 superficial layer at a depth of 220 um, corresponding to that marked by the red box in the top panel. Bars, 100 μm. (**b**) Mean fluorescence changes (ΔF/F) evoked by three pure tone stimuli (0.8, 1.0, and 1.2 KHz, duration 0.2 s, 8 trials each) for 4 example cells (marked by red circles in **a**). Scale bar, 0.5s and 20% ΔF/F. (**c**) ΔF/F with time for 20 pure tone-responsive cells during sequential step-wise application of pure tones from 0.5 to 16 kHz (black bar, tone duration s), with the magnitude of ΔF/F color-coded by the continuous scale shown on the right. Red trace on top shows the averaged ΔF/F for all 20 cells. Note that the BF for the vast majority of cells was located at 1.2 KHz. Scale bar, 0.5s and 20% ΔF/F. (**d**) Spatial distribution of all tone-selective cells in the example imaging plane as that shown in **a** (bottom), including data recorded at three different sound levels (60, 70, and 80 dB) of the same frequency. (**e**) 3D distribution of A1 cells with similar BFs at three different cortical depths, with sound stimuli of three different intensities applied.

## Discussion

Using *in vivo* two-photon calcium imaging to monitor the activities of a large population of A1 neurons in responses to pure tones of different frequencies, we found that the frequency preference of marmoset A1 neurons is homogeneous over distances of hundreds of micrometers in both horizontal and vertical directions relative to the pia surface. This was shown by using either acute loading of fluorescence Ca^2+^ indicator Cal-520 or *in vivo* expression of genetically encoded Ca^2+^ indicator GCaMP6f. These results demonstrated the feasibility of simultaneous recording of large populations of cortical neurons in the marmoset brain at single-cell resolution. Furthermore, macroscopic tonotopic maps previously observed by electrophysiological recording^7^ and intrinsic optical signals^8^ directly reflect homogeneous neuronal tuning properties within local regions of marmoset A1.

The precision of tonotopic organization at the cellular level has recently become a controversial issue^33–37^, based mostly on studies from mice. Although tonotopic map of monkey A1 at the cellular level has not been reported, previous studies using extracellular electrophysiological recording of neuronal activities have suggested that the tonotopic preference of cortical cells within local regions of A1 is homogeneous, consistent with a smooth tonotopic organization. Since both electrophysiological recording and intrinsic optical imaging have a spatial resolution of 50-100 μm^7,8,38^, the smooth tonotopic maps may result from the averaged responses over many neurons. Extracellular recordings could also be biased toward highly active neurons in either multi- or single-unit recordings^39^. *In vivo* two-photon calcium imaging approach offers cellular resolution that could unequivocally address the issue of local homogeneity in the frequency tuning properties of A1 neurons.

Several recent studies^9,10,40,41^ using the *in vivo* two-photon imaging approach have cast doubt on the existence of strictly tonotopic maps in A1. It was found that neurons in mice A1 with diverse frequency preferences were highly mixed locally, despite the presence of apparent macroscopic tonotopy. These previous studies revealed that the best frequency (BF) for evoking neuronal responses was highly variable among neurons within 50-100 μm distances, with differences in BFs as large as 2-4 octaves^42^. In the present study of marmosets, we found that differences of BFs among cells within 250 μm were less than 1 octave. To ensure that the discrepancy between our results and previous findings on mice was not caused by the difference in the Ca^2+^ imaging method, we also examined the local tonotopic property of rat A1 cells using our Cal-520 loading method. Quantitative comparison of our marmoset and rat results showed that rat A1 cells with different BFs were distributed in a much more mixed manner than that found in the marmoset A1, implicating different A1 organizations in rodents vs. marmosets. Nevertheless, in both rat and marmoset A1, cells with the same BFs were distributed in a manner that was far from random. Furthermore, we found that in each local rat A1 region, there was a large majority of cells with a particular BF, with other cells of diverse BFs inter-dispersed among them. The presence of a dominant population of cells with the same BF could account for the global tonotopic maps found previously in rodents^6,26^ and in our rat results (**Supplementary Fig. 4**), as well as the previously reported “clustered” and “salt-and-pepper” distributions of cells in different local regions in mouse A1^41,42^.

In highly visual animals such as monkeys and cats, neurons with similar receptive field properties in the primary visual cortex (V1) are well organized into local columns. By contrast, in rodents with poor vision, neurons with different receptive field properties are mixed locally in a “salt-and-pepper” manner^21,22,43–45^. By analogy, highly uniform tuning properties of A1 neurons locally in marmosets may reflect an organization principle favorable for auditory processing in animal species that are rich in vocal communication and social interaction. Natural sounds with syllabuses in marmoset calls need to be first decomposed into frequency-specific signals and then re-integrated for auditory perception. Previous studies have shown that some neurons in marmoset A1 selectively respond to the “twitter” sound syllabus, but not to the time-reversed one^46,47^. Integration of frequency-specific signals into syllabus may depend on intracortical connections among A1 neurons, in addition to potential contributions from thalamocortical inputs. Given the local homogeneity in the frequency tuning of A1 neurons of marmoset, long-range intracortical circuitry among neurons in different tonotopic A1 domains may play a significant role in the integration of auditory signals of different frequencies. The current results thus pave the way for further analysis of A1 circuitry underlying natural sound processing in marmosets.

## Acknowledgments

We thank Yang Dan, Ninglong Xu, and Siyu Zhang for suggestions and comments on the manuscript. We thank Neng Gong, Hao Li, and Xuebo Li for technical support. This work was supported by grants from Strategic Priority Research Program of Chinese Academy of Science, Grant No.XDBS0100000 and Shanghai Municipal Government Bureau of Science and Technology (16Jc1420200). The authors declare no competing interests.

## Author Contributions

H.H.Z., Z.M.S. and M.M.P. designed the research; H.H.Z. and J.F.H. performed all experiments with contributions from M.C. and Y.Q.W.; H.H.Z. analyzed the data; H.H.Z., Z.M.S. and M.M.P. wrote the manuscript.

## METHODS

### Animals

Animal care and experimental procedures were approved by the Animal Care Committee of Shanghai Institutes for Biological Sciences, Chinese Academy of Sciences (Shanghai, China). Four adult common marmosets (*Callithrix jacchus*; 1 male and 3 females; body weight: 260-400g) obtained from the non-human primate facility of the Institute of Neuroscience were used in this study. The marmosets were individually housed in a temperature- and humidity-controlled facility (26°C-30°C, 12 h light/dark cycles), and supplied with *ad libitum* water and balanced diet.

### Surgery

Prior to the surgery, the marmoset was anesthetized with a fentanyl cocktail (0.0005 mg/kg fentanyl citrate, 0.5 mg/kg midazolam and 0.05 mg/kg dexmedetomidine, all administered intramuscularly or intraperitoneally). After induction, anesthesia was maintained by 1.5%-3% isoflurane with pure oxygen, and animal was kept in a customized apparatus in a prone position. The body temperature was maintained at ∼37°C using a heating blanket and monitored with a rectal thermal probe. During the surgery, the anesthesia state was confirmed by the absence of pinch-induced tail or paw reflexes. A titanium head-post was attached to the parietal bone by dental cement, performed under sterile conditions. An 8 or 10 mm-diameter circular craniotomy and durotomy were performed to expose the auditory cortex, and a custom-made chronic window was implanted over the craniotomy and sealed with dental cement. The chronic window was made by fitting a coverslip (8-10 mm in diameter and 0.17–0.2 mm in thickness) to a titanium ring, and glued with silicone adhesive (KN-300X, Kanglibang, China). Animals were allowed to recover for at least 7 days. Antibiotics were intramuscularly administered for 3 consecutive days after surgery.

In one marmoset, we performed injection of GCaMP6f viral vector, using a procedure previously described^1^. In brief, after anesthesia was induced as described above, the chronic window was carefully removed. A pulled glass pipette (30 μm outer-diameter) loaded with 3-4 μl virus solution was inserted into the auditory cortex at a depth of ∼500 μm. The viral preparations were adjusted to the final concentration of 1-3×10^12^ vg ml^-1^ for rAAV2/9-hSyn-tTA and 2-8×10^12^ vg ml^-1^ for rAAV2/9-TRE3-GCaMP6f and 1-2 μl of equal-volume mixture of these AAVs was injected within 10-15 min. A new chronic window was pressed onto the brain surface and attached to the skull by the dental cement. Two-photon imaging began ∼ 3 weeks after injection.

### Imaging of intrinsic optical signals

Marmosets were anesthetized by 1.5%-2% isoflurane with pure oxygen through a custom-made mask. The head was immobilized with a customized apparatus. During imaging sessions, the anesthesia was switched to fentanyl cocktail (0.001 mg/kg fentanyl citrate, 1 mg/kg midazolam and 0.1 mg/kg dexmedetomidine, administered intraperitoneally). In our hands, the marmosets remained in a stable state for at least 3 hours, as indicated by the heart rate, blood oximetry, and in some cases electroencephalogram. In some cases, we applied another fentanyl cocktail injection for prolonged imaging sessions. Signals of reflectance change (intrinsic hemodynamic signals) corresponding to the local cortical activity were acquired (Imager 3001, Optical Imaging Inc., Germantown, NY) with 660-nm illumination. Signal-to-noise ratio was enhanced by trial averaging (15–30 trials per stimulus condition). Acoustic stimuli were presented in blocks. Each block contained all stimulus conditions (e.g., various frequencies) and a blank condition. For each condition, imaging began 0.6 s before the sound stimulus onset (for baseline signals). The total imaging time for each condition was 6 s, during which 30 consecutive frames were collected (i.e., 5 Hz frame rate). All stimulus conditions were displayed in a randomized order.

### Acoustic stimuli

All acoustic stimuli were generated using MATLAB (MathWorks) and saved uncompressed (96-kHz sampling frequency, 16 bits). The sound delivery system was calibrated using B&K (2669-L) calibrator. The sound stimuli were presented by a Fostex FT28D speaker (0.5-32 KHz) or KEF ls50 speaker (used in most cases, 1-16 KHz), placed 30 cm (15 cm in two-photon imaging) to the contralateral ear. For imaging intrinsic signals, pure tone stimuli (4 clicks, 0.2 s duration and 0.5 s interval, 2.3s in total) were used. For two-photon imaging, the duration of pure-tone stimuli was 0.2 s (including 5-ms ON and OFF linear ramps), with inter-stimulus interval of 1 to 1.5 s. Each combination of frequency and intensity was presented 5 or 8 times. The intensities used in experiments ranged from 30 to 90 dB SPL in 10 dB increments. Our customized sound-insulating chamber can attenuate most ambient noise (< 30 dB SPL in the chamber).

### In vivo two-photon calcium imaging

Marmosets were anesthetized as described above in optical imaging experiments. The dental cement around the chronic window was removed and replaced with a new chronic window with a pin hole (i.e.,∼ 500 μm in diameter), which allowed glass micropipette access to the cortex for dye loading. The new chronic window was immobilized by dental cement. A cell-permeant calcium indicator Cal-520 AM (AAT Bioquest) was injected into layer 2/3 auditory cortex as previously described^2^. In brief, Cal-520 AM (40-50 μg) was dissolved with 4 μl Pluronic/DMSO mixture (20% w/v Pluronic F-127 in dimethyl sulfoxide) and the dye solution was diluted with 1 μl of a red dye (Alexa Fluor 594, 2 mM stock solution) and 35 μl of pipette solution (in mM: 10 HEPES, 2.5 KCl, 150 NaCl, pH 7.4.). The dye solution was sonicated and filtered with a 0.22 μm filter to remove dye aggregates, and then transferred to a glass pipette (2-4 μm tip). Dye ejection was performed in layer 2/3 of the auditory cortex (200-300μm from the surface) at 6-12 psi for 60-90 pulses at 1Hz using Picospritzer (Parker Hannifin, Hollis, NH, USA). Two-photon calcium imaging was performed 1-2 hours after successful dye ejection. Fluorescence from neurons was monitored with a custom-made LotosScan microscope (LotosScan, Suzhou Institute of Biomedical Engineering and Technology) and coupled with a mode-locked Ti:Sa laser (model “Mai-Tai DeepSee”, Spectra Physics). The excitation wavelength was fixed at 920 nm. Imaging was performed using a 16X, 0.8 NA objective (Nikon) or a 40X, 0.8 NA objective (Nikon). Technical limitations of the microscope kept us from overfilling the back aperture of the 16X objective, which may reduce the effective NA. The beam size was large enough to overfill the back aperture of the 40X objective. Images were acquired at a frame rate of 40 Hz.

### Rat experiments

Three Sprague-Dawley rats (female; body weight: 260-400g) were used in the experiments. For rat experiments, anesthesia was maintained by 1.5%-3% isoflurane with pure oxygen and body temperature was maintained at 36–38 °C. Surgery procedure was similar with that in marmoset. The primary auditory cortex was located as described in previous studies^3,4^ by combining stereotactic coordinates and intrinsic optical imaging (see Supplementary figure 4). Other experimental procedure was the same with marmoset. During functional imaging experiments, the anesthesia was switched to fentanyl cocktail (0.025 mg/kg fentanyl citrate, 2.5 mg/kg midazolam and 0.125 mg/kg dexmedetomidine, administered intraperitoneally).

### Data analysis

Images were analyzed in Matlab (Mathworks) and ImageJ (National Institutes of Health). For correcting lateral motion in the imaging data, a rigid-body transformation based frame-by-frame alignment was applied by using Turboreg plugin (ImageJ software). Cells were identified by hand on the basis of size, shape, and brightness. Fluorescence changes with time of individual cell were extracted by averaging pixel intensity values within cell masks in each frame. Data were discarded if brain pulsation were evident during imaging. Fluorescence intensity change (ΔF_t_) evoked by each stimulus (t = 0.2 s + 0.2 s; stimulus duration, 0.2 s; post-stimulus signal 0.2 s) were normalized by the pre-stimulus baseline fluorescence (F_0_, 0.5 s). For each stimulus, the mean change in fluorescence (ΔF/F) was calculated by averaging responses to all trials for each stimulus condition. Cells showing significant differences in the fluorescence intensity signals observed during baseline vs. stimulus-presentation periods (P<0.05, ANOVA) were defined as “responsive cells”. Of these, tone-selective cells were defined by significant differences in intensity responses signals across all frequencies (P<0.05, ANOVA). The “best frequency (BF)” is defined as the frequency of sound stimulus that evoked the highest response. In our data set, nearly all responsive cells were selective to pure tones. Error bars indicate SEM. In cellular tonotopic maps, only tone-selective cells were color coded and analyzed. In Fig. 5 for rat data, tone-selective cells that at any sound level were used to obtain the tonotopic maps.

Frequency vector maps were calculated using the vector-summation method^5^. Single-condition maps were averaged and spatially smoothed (Gaussian, σ = 2 μm). Fluorescence change (ΔF) maps were obtained by subtracting baseline images from images of responses evoked by sound of one of six frequencies. For each pixel, a vector is calculated based on its response to six different frequencies (vector summation). The angle represents the frequency preference at each pixel (angles of the vectors were color-coded) and the magnitude represents the strength of the frequency-selectivity at each pixel (length of the vectors coded by color intensity).

